# 17β-Estradiol and Estrogen Receptor α Promote Right Ventricle Angiogenesis in Pulmonary Hypertension via Apelin Signaling

**DOI:** 10.64898/2025.12.11.693201

**Authors:** Simon Bousseau, Andrea L. Frump, Avram Walts, Amanda Fisher, Bakhtiyor Yakubov, Steeve Provencher, Sandra Breuils Bonnet, Sébastien Bonnet, Tim Lahm

**Affiliations:** National Jewish Health, Denver, CO, USA; Indiana University School of Medicine, Indianapolis, IN, USA; Laval University, Québec City, QC, Canada; University of Colorado, Anschutz Medical Campus, Aurora, CO, USA; Rocky Mountain Regional VA Medical Center, Aurora, CO, USA

**Keywords:** Endothelial cell function, right heart failure, sex hormones, angiogenesis

## Abstract

Right ventricular (RV) failure is the major cause of mortality in pulmonary hypertension (PH). Adaptive angiogenesis and RV endothelial cell (RVEC) function are major modifiers of RV adaptation in PH, but the underlying mechanisms and their regulators remain incompletely understood. RV adaptation in PH is sexually dimorphic, and 17β-estradiol (E2) exerts protective effects on RV cardiomyocytes. Whether E2 modifies angiogenesis and RVEC function in RV failure remains unknown.

We hypothesized that E2 and estrogen receptor α (ERα) promote RV angiogenesis and RVEC homeostasis in PH and aimed to identify underlying mechanisms.

We assessed E2’s angiogenic effects using cultured human cardiac microvascular endothelial cells (hCMVECs), RVECs from PH patients with RV failure, and RVECs from sugen/hypoxia (SuHx) and monocrotaline (MCT) rat models. *In vivo*, we evaluated RV capillary density in PH rats treated with E2 or ERα-selective agonist. Apelin signaling was evaluated via apelin receptor blockade.

E2 enhanced angiogenesis in male hCMVECs and RV capillary density in female SuHx-PH rats. E2 reversed angiogenic alterations in RVECs from SuHx-PH rats via apelin receptor signaling. In RVECs from PH patients with RV failure, E2 stimulated vascular network formation. In rat and human PH-RVECs, ERα was necessary and sufficient to mediate E2-induced angiogenesis. Activation of ERα with ERα-specific agonist restored RV capillary density in vivo. ERα-mediated angiogenesis required apelin signaling.

These data indicate that E2 promotes RV angiogenesis via ERα and apelin signaling and identify a novel ERα–apelin axis in RVECs as a potential therapeutic target to restore RV vascular integrity in PH.

## Introduction

Depressed right ventricle (RV) function is a common and strong predictor of mortality across different groups of pulmonary hypertension (PH) ^1,2^. As RV afterload increases during the development of PH, the RV exhibits compensatory mechanisms that include structural changes, neurohormonal activation, and increased contractility^3–5^. On a cellular level, these changes are accompanied by alterations in angiogenesis, changes in mitochondrial function and substrate utilization, increased reactive oxygen species, changes in myosin isoform expression, and altered sarcomere organization^4,5^. This allows for a state of adaptive RV hypertrophy (RVH), characterized by a cardiac output that is still sufficient to meet the metabolic demands of the body^3–6^. However, once the RV’s compensatory mechanisms are exhausted, the RV transitions from adaptive (compensated) to maladaptive (decompensated) RVH, and RV failure (RVF)^3–6^. There is an unmet need in PH to prevent, delay or reverse the transition from adaptive to maladaptive RVH^3–6^

Angiogenesis has recently emerged as a major modifier of RV adaptation in PH^7^. Adequate and well-organized angiogenesis is critical to maintaining substrate delivery to the hypertrophied myocardium^7,8^. In addition, the endothelium serves as a paracrine “machine” that regulates the function of the surrounding myocardium^3,9^. Insufficient and dysregulated angiogenesis and microvascular ischemia are purported critical contributors to RVF development^8,10^. In fact, lack of coordinated angiogenesis has been identified as a critical mediator of the transition from adaptive to maladaptive RVH^7^. However, the mechanisms responsible for RV endothelial cell (RVEC) dysfunction, angiogenesis dysregulation, and microvascular ischemia are not understood.

Importantly, RV adaptation in PH is sexually dimorphic, and female PH patients exhibit better RV function than their male counterparts^11,12^. In multiple groups of PH, female sex is associated with better survival^1,11,13^, and this survival advantage has been linked to better RV function^11^. Several lines of evidence demonstrate that female sex hormones contribute this sexual dimorphism in RV function. In healthy postmenopausal hormone therapy users, higher plasma levels of 17β-estradiol (E2; the most abundant female sex steroid) are associated with increased RV ejection fraction^14^. We have previously identified several molecular mechanisms of how E2 affects RV function in PH. In particular, we reported that 1) E2 attenuates PH-induced decreases in cardiac output and exercise capacity^15^, 2) E2 favorably affects several pathogenetically relevant processes in the failing RV, such as pro-apoptotic signaling, pro-inflammatory cytokine activation, and mitochondrial dysfunction^15^, 3) E2 attenuates RVF induced by acute strenuous exercise^16^, 4) E2 improves autophagic flux and attenuates pro-apoptotic signaling in exercise-induced RV falure^16^, and 5) E2 employs and activates a novel estrogen receptor (ER) α-BMPR2-apelin axis that mediates RV adaptation *in vivo* and promotes RV cardiomyocyte resilience as well as beneficial RV cardiomyocyte - RVEC interactions *in vitro*^17^.

E2 signals *via* its two main receptors ERα and ERβ^18^. We previously demonstrated that 1) ERα is localized in human RV cardiomyocytes and RVECs^17^, 2) treatment with E2 attenuates PH-induced decreases in RV ERα expression^15^, 3) ERα-selective agonist treatment recapitulates E2’s protective effects on anti-apoptotic and pro-angiogenic signaling^15,17^, 4) ERα is necessary for E2’s RV-protection^17^, and 5) selectively activating ERα is sufficient to elucidate RV protection^15,17^. Given ERα’s location in RVECs, and given ERα’s known salutary effects on endothelial cells in various organ systems^8,19^, we now sought to determine whether the E2-ERα axis is active in RVECs. We hypothesized that E2, via ERα, exerts protective effects on RVEC angiogenic function in health and RVF and improves vascularization in animal models of RVF. We demonstrate that E2 stimulates angiogenesis in RVECs from PH patients with RVF and enhances RV vascularization in PH and RVF models through ERα and apelin.

## Materials and Methods

(Please see the data supplement for detailed methods)

### In vivo studies

Animal models employed were sugen/hypoxia-induced PH (SuHx-PH) and monocrotaline-induced PH in male and female rats as described previously^15,17^. Subgroups of animals were treated with E2 or modulators of ER or apelin receptor signaling as described previously^15,17^. Subgroups of females were ovariectomized (OVX) as described previously^15,17^. Vascularization was assessed via lectin staining and capillary density quantification. Cardiopulmonary hemodynamics, and RV structural, functional and molecular alterations of the animals employed were previously published^15,17^.

### RVEC isolation and culture

*Rat RVECs:* RVs were dissected from male normoxic or SuHx-PH Sprague-Dawley rats, minced and digested using Collagenase II (Gibco, ThermoFisher; Waltham, MA) at 37°C for 1-3 hrs as previously described. Cells were selected using Pan-mouse IgG Dynabeads (ThermoFisher) coated with mouse monoclonal anti-rat CD31 antibody (BD Biosciences; BD Biosciences) and then seeded on a gelatin-coated 6-well plate in EGM-2MV (Lonza; Basel, Switzerland) phenol red-free media with charcoal stripped FBS (ThermoFisher) substituting regular FBS and supplemented with Normocin (Invivogen, San Diego, CA). Endothelial lineage was validated based on morphology, endothelial marker expression, pseudo-vascular network formation, and Dio-AC-LDL uptake. Cells were utilized in experiments up to passage 7. To study E2 effects in an E2-naïve background, we focused on cells from male animals.

*Human RVECs:* RVECs from PH patients with RV failure (PH-RVECs) were isolated as previously described^23^. Patient characteristics are provided in **table 1**. Cells were grown in phenol red-free Endothelial Cell Growth Basal Medium-2 supplemented with 0.1% human epidermal growth factor, 0.1% ascorbic acid, 0.1% GA-1000 (gentamicin, amphotericin B), 2% fetal bovine serum, 0.1% VEGF, 0.4% human basic fibroblast growth factor, 0.04% hydrocortisone, 0.1% R3-Insulin-like growth factor −1, and 1% streptomycin-penicillin (all chemicals were purchased from Lonza). Cells were grown at 37°C in a humidified incubator gassed with 5% CO_2_ and were used for experiments at the third to ninth passage in charcoal stripped media.

**Table 1:** PH Patient characteristics.

| **Tissue** | **Patient Sample Name** | **Group** | **PH diagnostic** | **Sex** | **Age** | **RV dilated?** | **RV function** | **Tissue Source** |
| --- | --- | --- | --- | --- | --- | --- | --- | --- |
| RV | 1 | Decompensated RV failure | Limited SSc-PAH | F | 47 | RV severely dilated | Severely reduced | Autopsy |
| RV | 2 | Decompensated RV failure | Heritable PAH | F | 61 | RV severely dilated | Severely reduced | Autopsy |
| RV | 3 | Decompensated RV failure | COPD | M | 57 | RV severely dilated | N/A | Autopsy |

*Human cardiac microvascular ECs* (hCMVECs; of male origin) were purchased from Lonza and grown and maintained in EGM-2MV media as described for human RVECs. Cells were utilized in experiments up to passage 9.

Functional endpoints assessed were cell proliferation, vascular network formation, and migration. Molecular endpoints were assessed according to standard methods.

## Results

### E2 increases angiogenic capacity and angiogenesis mediator expression in cultured hCMVECs

We first assessed effects of E2 on hCMVEC angiogenic capacity *in vitro*. We found that E2 indeed increased hCMVEC proliferation, migration and vascular network formation (**Fig. 1A-E**). This was accompanied by an increase in expression of the pro-angiogenic factors VEGFR2, p-eNOS (Ser 1177), and apelin **(Fig. 1F-H)**. Together, these results indicate that E2 dose-dependently increases hCMVEC angiogenic capacity and expression of pro-angiogenic mediators.

**Fig. 1.**
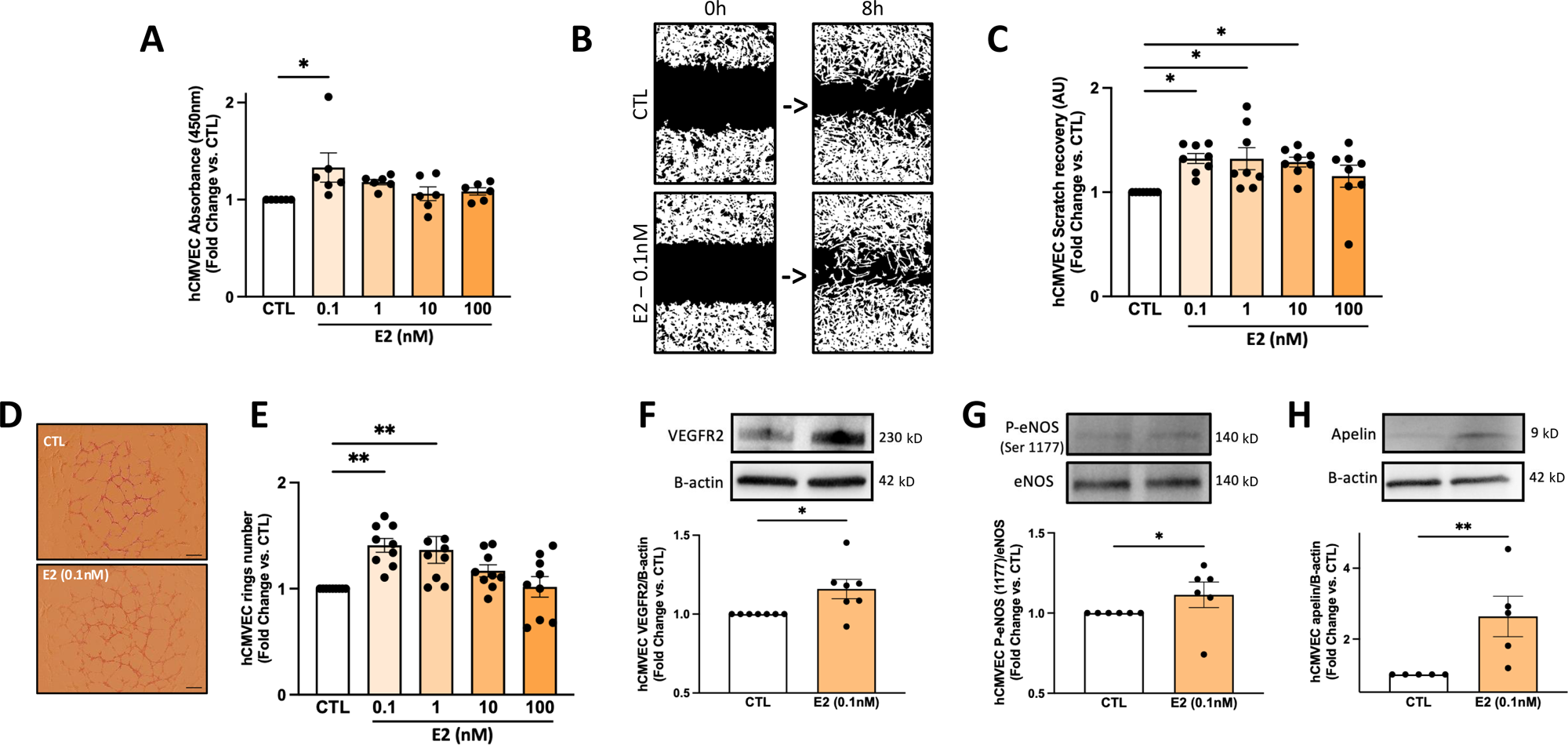
E2 increases *in vitro* angiogenesis in male hCMVECs. **A)** Proliferation assessed by cell counting kit-8. **B, C)** Migration assessed via wound healing assay (representative images shown in B; quantification shown in C). **D, E)** Vascular network (ring) formation assessed by matrigel assay (representative images shown in D; quantification shown in E) Images are 4x; scale bar is 100 μm. **F)** VEGFR2 expression, **G)** eNOS activation (eNOS phosphorylated at Ser1177 normalized for total eNOS) and **H)** Apelin expression by Western blot (representative images and densitometric quantification). Concentrations are as indicated in figure; duration of treatment was 24 hours. Each data point represents one independent experiment. Error bars are means±SEM. *p<0.05, **p<0.01 vs CTL by ANOVA (A-E) or t-test (F-H).

### E2 increases RV capillary density in vivo

While E2 signaling stimulates tube formation in hCMVECs **(Fig. 1D-E)**, exerts RV-protective *effects in vivo*, and increases RV expression of the pro-angiogenic peptide apelin ^17^, it remains unknown whether E2 attenuates PH-induced capillary rarefaction. We therefore assessed RV capillary density using isolectin staining in female intact and ovariectomized (OVX) SuHx-PH rat RVs (hemodynamics published in ^15^). In this cohort, female SuHx-PH rats exhibited a decrease in RV cardiac index as compared to healthy controls^15^. This was further reduced by OVX^15^. On the other hand, E2-replete OVX SuHx-PH rats maintained RV adaptation^15^.

Intact SuHx-PH rats exhibited a 44% reduction in capillaries per field compared to controls **(Fig. 2A-B)**. This loss of vascular integrity was exacerbated in SuHx-PH OVX rats, which exhibited a 53% reduction in capillaries per field vs normoxic rats and a modest but significant reduction vs intact SuHx-PH rats **(Fig. 2A-B)**. This suggests a more profound decrease in the functional capillary network in the SuHx-PH RV in absence of endogenous sex hormones. Building on this observation, we found that SuHx-PH OVX females replete with E2 exhibited a 45% increase in capillaries per field as compared to OVX SuHx-PH females without E2 **(Fig. 2A-B)**. These data, which parallel E2 effects on RV function^15^, suggest that E2 exerts pro-angiogenic effects in SuHx-PH-induced RVF and that E2’s protective effects against RVF may be at least in part mediated through maintenance of the RV capillary network.

**Fig. 2:**
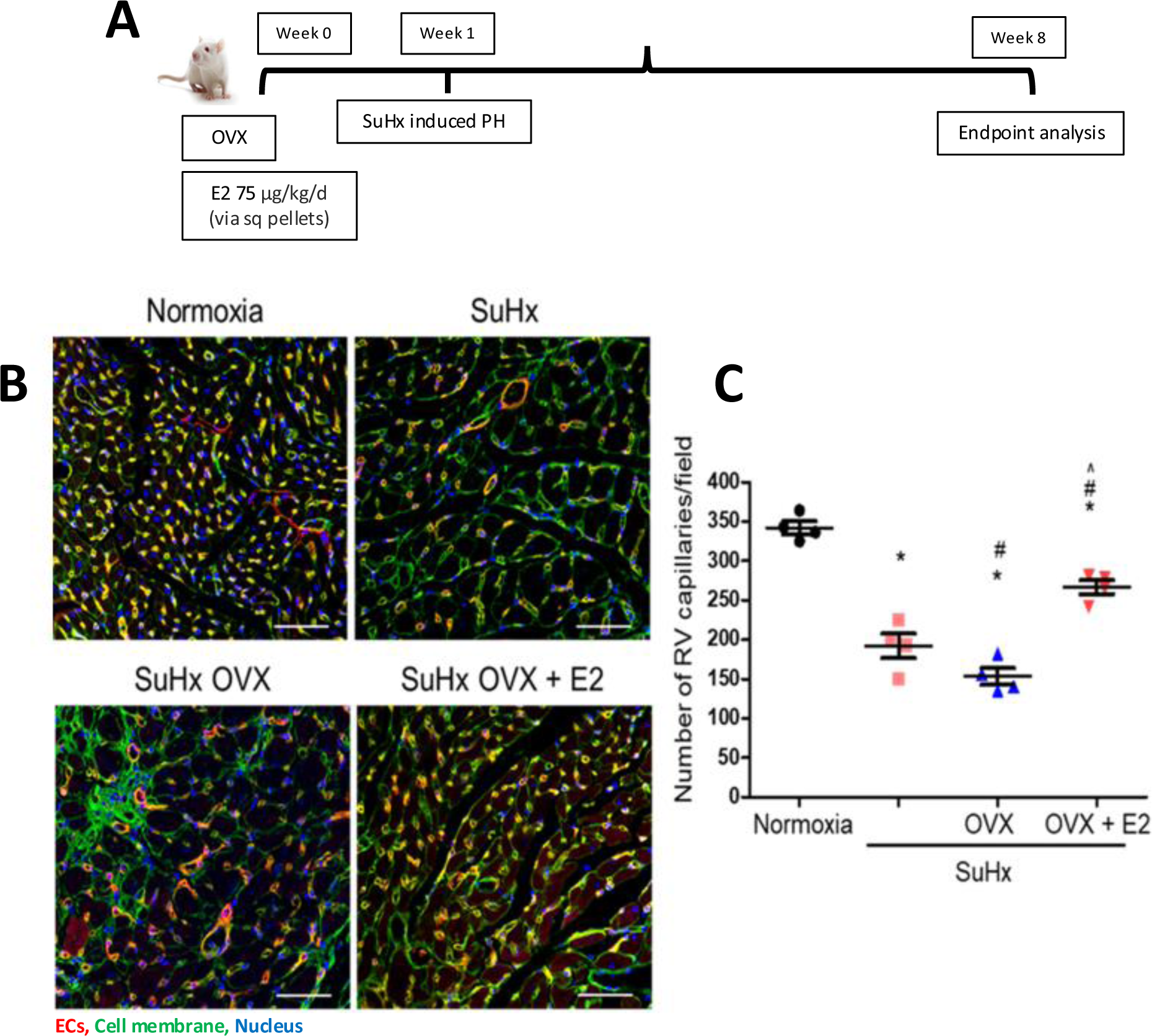
E2 increases RV capillary density in female sugen/hypoxia (SuHx)-PH rats. **A)** Experimental design. OVX = ovariectomy. **B)** Immunofluorescence overlay of lectin Griffonia simplicifolia (red, endothelial cells (ECs)), wheat germ agglutinin (green, cell membrane) and DAPI (blue, nuclei). Scale bar=50 μm. **C)** Quantification of capillary density as RV capillaries/field. Error bars are means±SEM. *p<0.05 vs normoxia, ^#^p<0.05 vs SuHx, ^p<0.05 vs SuHx OVX untreated, by ANOVA.

### E2 attenuates PH-induced impairments in RVEC angiogenic function

To delineate effects of E2 directly on RVEC angiogenesis, we isolated RVECs from SuHx-PH rats treated with E2 in vivo. In this experiment, we employed male normoxic control, untreated SuHx-PH rats and SuHx-PH rats treated with E2. We performed tube formation assays and found that RVECs isolated from SuHx-PH rats exhibited fewer vascular networks compared to RVECs isolated from normoxic controls **(Suppl. Fig. 1)**. Meanwhile, RVECs isolated from SuHx-PH rats treated with E2 formed significantly more networks compared to RVECs from untreated SuHx-PH rats **(Suppl. Fig. 1)**. Together these data indicate that E2 directly targets RVECs and preserves angiogenic capacity in RVECs from male SuHx-PH rats.

### E2-mediated vascular network formation of cultured SuHx PH-RVECs is apelin receptor-dependent

In previous studies, we established the pro-angiogenic peptide apelin as a downstream target of E2 and identified that apelin is required for E2 to exert RV-protective effects^15,17^. We therefore evaluated the angiogenic effect of E2-apelin signaling in SuHx PH-RVECs. Specifically, we treated RVECs from male SuHx-PH rats with pyroglutamated apelin-13 (Pyr-Apelin). Pyr-Apelin indeed induced vascular network formation, indicating that apelin induces angiogenesis in SuHx PH-ECs **(Fig. 3)**. To determine whether apelin’s angiogenic effect is dependent on the apelin receptor (APLNR), SuHx PH-RVECs were co-treated with Pyr-Apelin + APLNR antagonist ML221. As expected, apelin’s effects on vascular network formation were attenuated after APLNR inhibition **(Fig. 3)**. We next assessed whether apelin signaling was necessary for E2-induced network formation. As previously shown, E2 induced network formation in SuHx PH-RVECs **(Fig. 3)**. However, when SuHx PH-RVECs were co-treated with E2 and ML221, E2-induced ring formation was attenuated **(Fig. 3)**. Taken together, these data indicate that E2-induced vascular network formation in SuHx PH-RVECs is dependent on intact APLNR signaling.

**Fig. 3:**
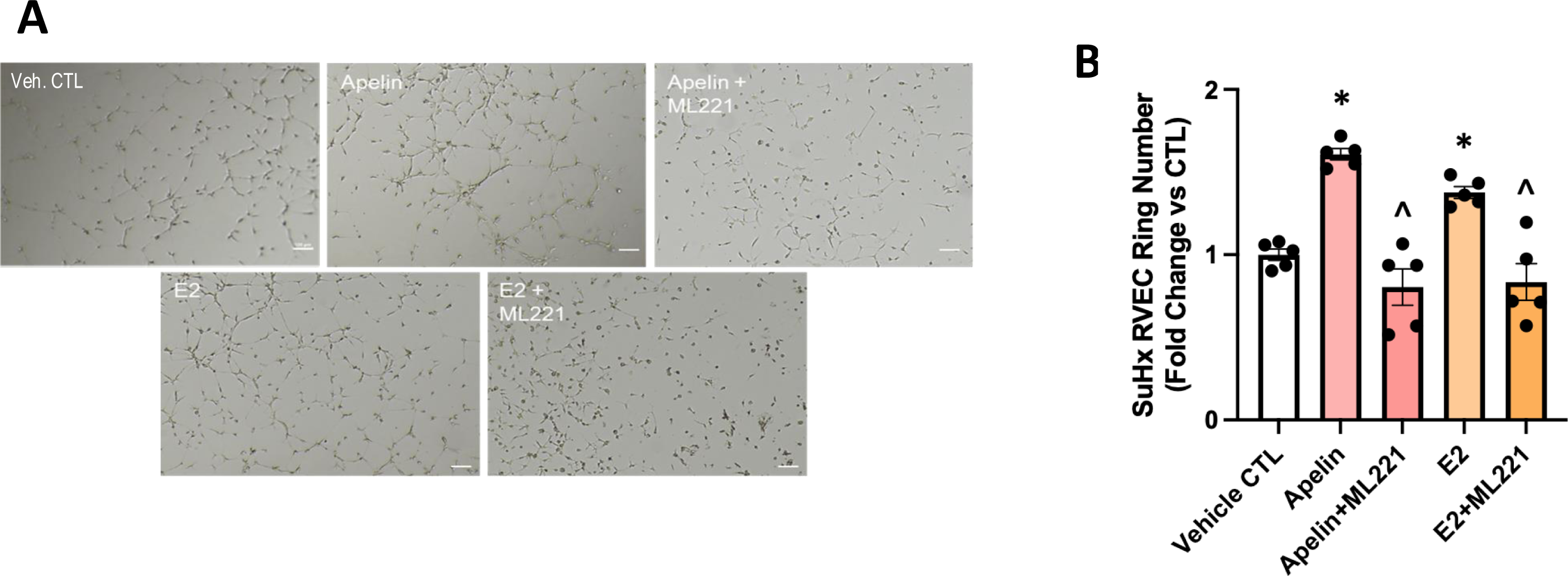
E2-mediated tube formation is apelin receptor-dependent in SuHx-PH RV endothelial cells (RVECs). **A) and B)** Matrigel tube formation assay using RVECs from male SuHx rats treated with vehicle, apelin (100 nM) or E2 (10 nM) ± apelin receptor antagonist ML221 (100 µM) *in vitro* for 24 hours (representative images in A and quantification in B). Images are 4x; scale bar is 100 μm. N=10 fields/condition. Error bars are means±SEM. Each datapoint = one animal. *p<0.05 vs vehicle CTL; ^p<0.05 vs apelin or E2 treatment respectively, by ANOVA.

### E2 exerts pro-angiogenic effects in RVECs from PH patients with RVF

After demonstrating that E2 increases angiogenesis in SuHx PH-RVECs, we sought to assess whether such effects are also observed in RVECs from patients with RV failure (PH-RVECs). While E2 did not affect human PH-RVEC proliferation **(Fig. 4A)**, it significantly increased migration and vascular network formation by ∼20% **(Fig. 4B-E)**. E2 also increased the abundance of the pro-angiogenic mediators VEGFR2, p-eNOS (Ser 1177) and angiopoietin-2 **(Fig. 4F-H)**. These findings demonstrate that E2 stimulates angiogenesis in cultured RVECs from PH patients with RVF.

**Fig. 4:**
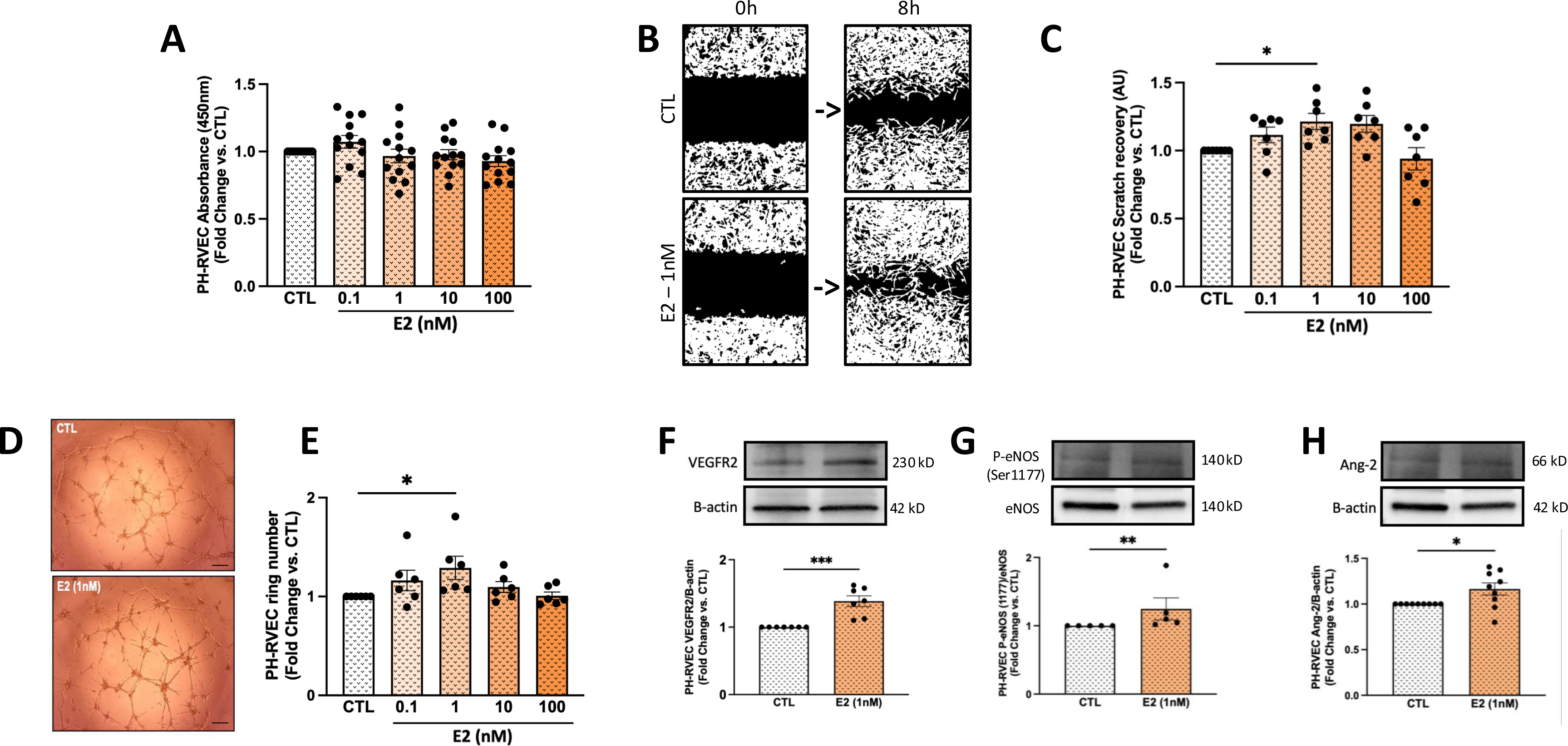
E2 stimulates angiogenesis in cultured RV endothelial cells (RVECs) from male and female PH patients with RV failure. **A)** Proliferation assessed by cell counting kit-8. **B, C)** Migration assessed via wound healing assay (representative images shown in B; quantification shown in C). **D, E)** Vascular network (ring) formation assessed by matrigel assay (representative images shown in D; quantification shown in E). Images are 4x; scale bar is 100 μm. **F)** VEGFR2 expression, **G)** eNOS activation (eNOS phosphorylated at Ser1177 normalized for total eNOS) and **H)** Angiopoietin-2 (Ang-2) expression by Western blot. Concentrations are as indicated in figure; duration of treatment was 24 hours. Each data point represents one independent experiment. Error bars are means±SEM. *p<0.05, **p<0.01 vs CTL by ANOVA (A-E) or t-test (F-H).

### E2 effects on angiogenesis are mediated via ERα and apelin

We next aimed to investigate through which of the two estrogen receptors is mediating E2’s pro-angiogenic effects. Utilizing RVECs from male SuHx-PH rats, we employed antagonists directed against ERα (MPP) or ERβ (PHTPP). ERα inhibition, but not ERβ inhibition, abrogated stimulatory effects of E2 on rat RVEC vascular network formation **(Suppl. Fig. 2)**, suggesting that ERα is necessary for E2 to exert pro-angiogenic effects. As shown previously, E2 also was unable to increase vascular network formation after APLNR inhibition.

In a next step, we sought to determine whether selective activation of ERα is sufficient to replicate pro-angiogenic effects of E2. Indeed, treatment of male rat RVECs with the highly ERα-selective agonist BTPα^17^ replicated stimulatory effects of E2 on vascular network formation **(Fig. 5)**. Interestingly, co-treatment with ML221 abrogated this effect, suggesting that APLNR is necessary for ERα to exert pro-angiogenic effects **(Fig. 5)**.

**Fig. 5:**
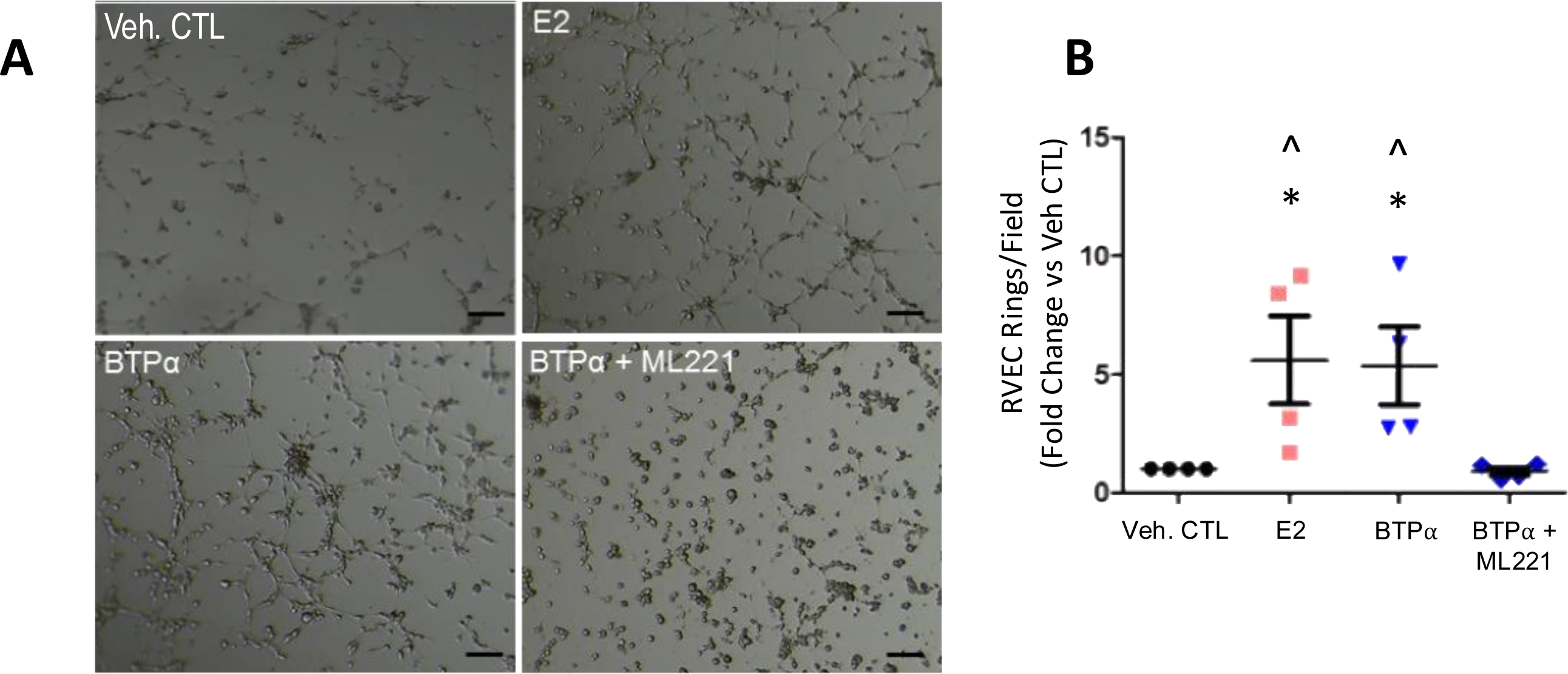
Pro-angiogenic effects of ERα in rat RV endothelial cells (RVECs) are apelin-dependent. **A) and B)** Vascular network (ring) formation assessed by matrigel assay using RVECs isolated from control male rats and treated with vehicle, E2, or BTPα (ERα agonist) ± APLNR antagonist ML221 *in vitro* (24 hours; representative images in A and quantification in B). Images are 4x. Scale bars = 100 μm. Each datapoint = one rat; 10 fields/condition. Error bars are means±SEM. *p<0.05 vs veh CTL; ^p<0.05 vs ML221 treatment by ANOVA.

Together, these data indicate that ERα is necessary for E2 to exert pro-angiogenic effects, that selective activation of ERα is sufficient to replicate pro-angiogenic effects of E2, and that ERα signals via the APLNR.

### Selective activation of ERα is sufficient to increase RV capillary density in vivo

Having established the effects of ERα *in vitro*, we next investigated whether ERα also exerts stimulatory effects on RV vascularization *in vivo*. To test this hypothesis, we first utilized the rat SuHx-PH model. A subset of male SuHx-PH rats was administered either ERα agonist PPT (850 ug/kg/day) or ERβ agonist DPN using a preventive approach (hemodynamics published in ^15^). E2-treated rats were employed as a positive control. Similar to female SuHx-PH OVX rats **(Fig. 2)**, male SuHx-PH rats exhibited a ∼50% decrease in capillaries per field compared to normoxic controls **(Fig. 6)**. On the other hand, treatment with E2 preserved the number of capillaries per field at levels similar to those of the control group. PPT-treated rats also demonstrated preserved capillary density compared to SuHx-PH alone (an increase of 25%), while DPN treatment was not associated with a significant increase in capillary density.

**Fig. 6:**
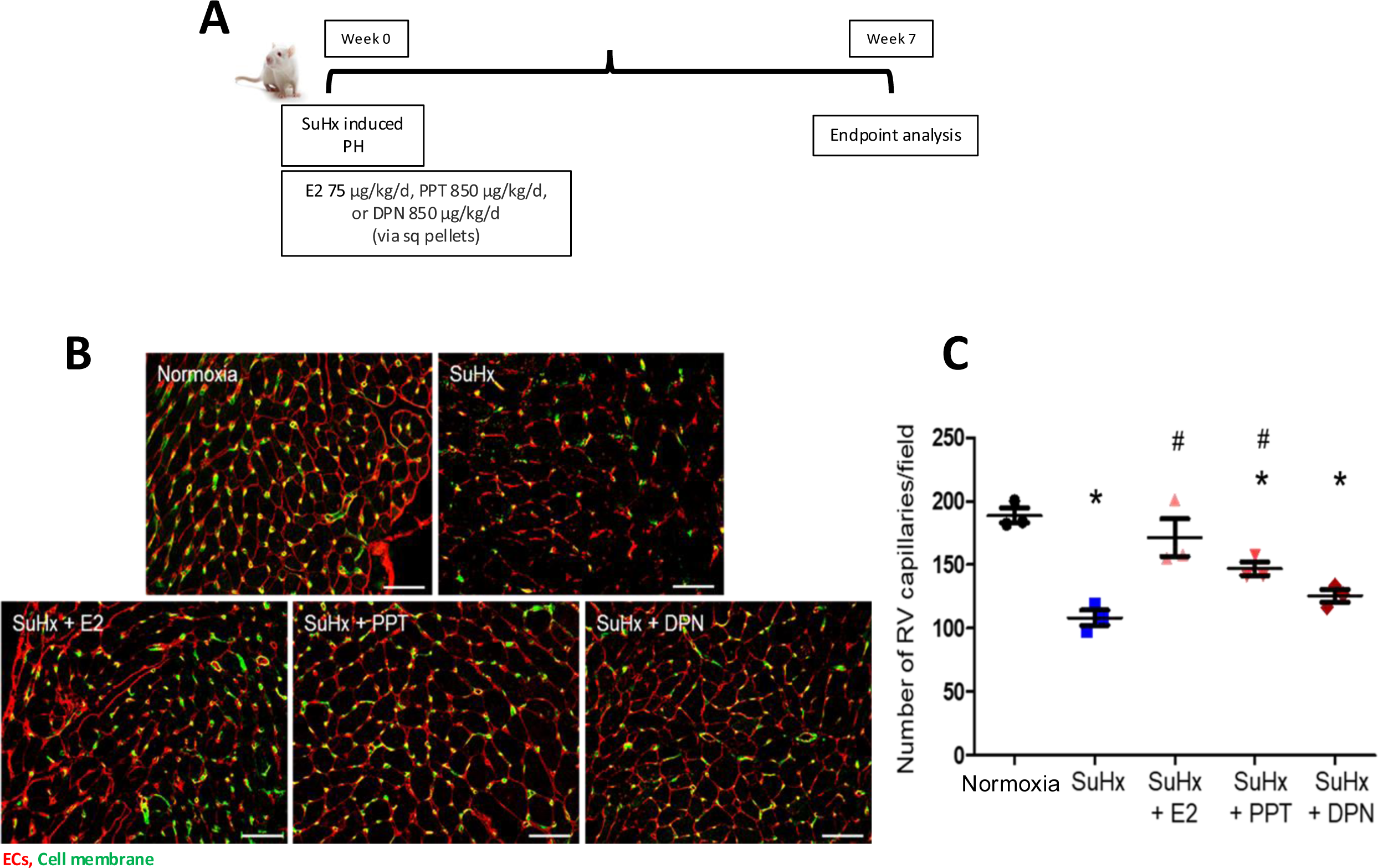
ERα activation with PPT is sufficient to prevent RV capillary dropout in male SuHx rats. **A)** Experimental design. **B)** Immunofluorescence overlay of lectin Griffonia simplicifolia (red, endothelial cells) and wheat germ agglutinin (green, cell membrane) in normoxic control, SuHx, SuHx+E2, SuHx+PPT and SuHx+DPN rats. Images are 20x. Scale bars=50 μm. **C)** Quantification of capillary density as RV capillaries/field. Each datapoint = one rat. Error bars are means±SEM. *p<0.05 vs normoxia; ^#^p<0.05 vs untreated SuHx by ANOVA.

Importantly, PPT was also able to *rescue* MCT-induced decreases in RV capillary density (again phenocopying E2 effects), when administered to male rats (hemodynamics published in ^17^; **Suppl. Fig. 3**). Taken together, these data indicate that E2 preserves and rescues RV capillary integrity in experimental RVF and that selectively activating ERα signaling replicates these effects.

### ERα is necessary and sufficient to promote tube formation in PH-RVECs

Lastly, we sought to determine if pro-angiogenic effects of ERα extend to human PH-RVECs. Indeed, silencing ERα in PH-RVECs resulted in an 80% decrease in vascular network formation **(Fig. 7A&B)**. In addition, E2’s effects on vascular network were abrogated by 60% after ERα silencing **(Fig. 7C)**. On the other hand, treating PH-RVECs with PPT replicated stimulatory effects of E2 on angiogenic capacity **(Fig. 7D)**. These data indicate that ERα indeed is necessary and sufficient to promote vascular network formation in RVECs from PH patients with RVF.

**Fig. 7:**
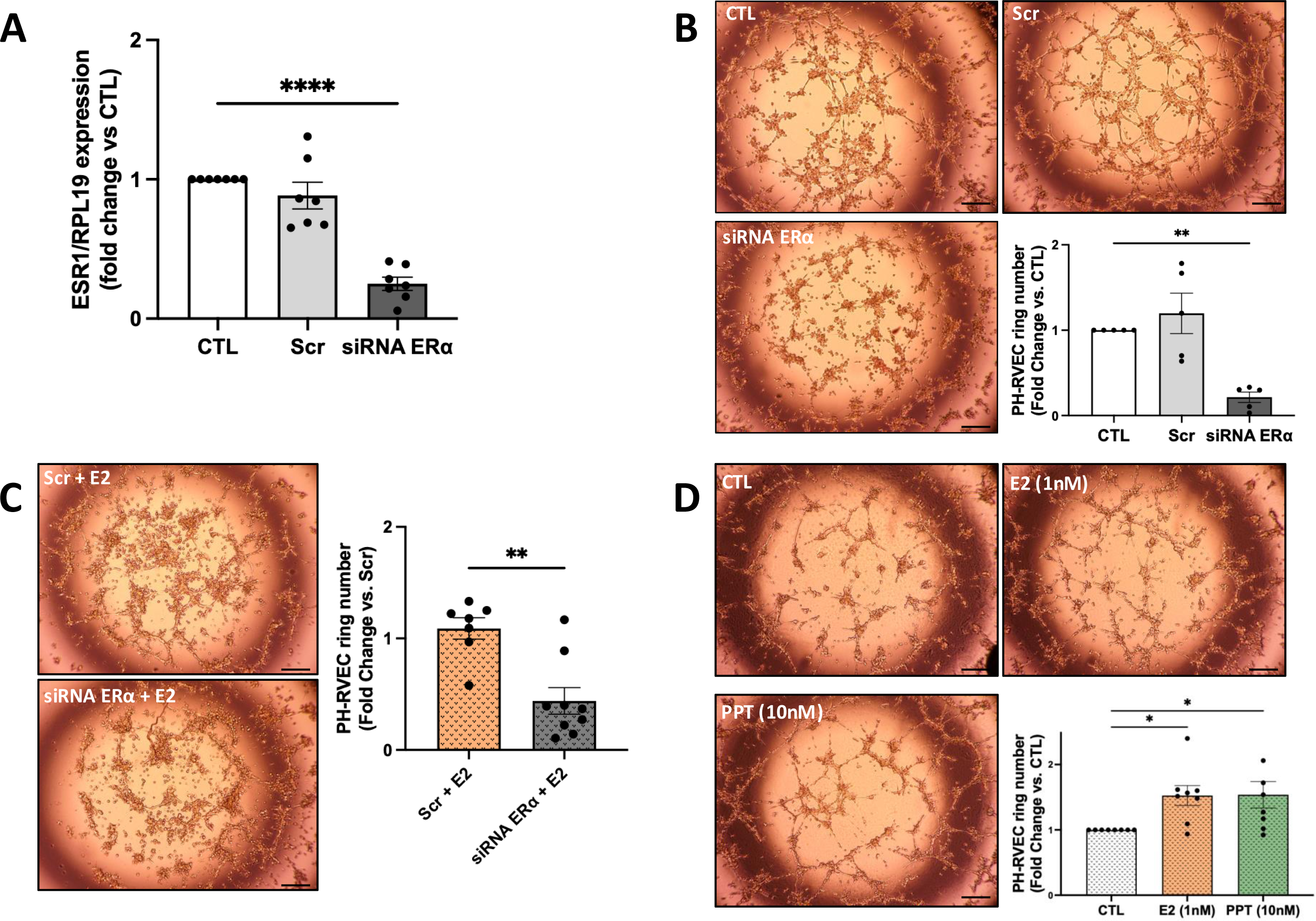
ERα is necessary and sufficient to promote tube formation in male and female PH RVECs. **A)** Evidence of successful ERα knockdown for 24 hours assessed by qPCR using RPL19 as a control. **B-C)** Representative images and quantification of RVEC vascular network formation by matrigel assay after siRNA knockdown in unstimulated conditions (B) and with E2 treatment (C). Note decrease in tube formation in unstimulated conditions (B) and with E2 treatment (C) after siRNA knockdown. CTL = untreated control; Scr = scramble control. **D)** Representative images and quantification of RVEC vascular formation by matrigel assay after treatment with E2 or ERα agonist PPT for 24 hours. Note phenocopying of E2 effects with PPT. Each data point is one independent experiment. Error bars are means±SEM. Images are 4x; scale bar is 100 μm. *p<0.05, **p<0.01, ****p<0.0001 vs CTL or Scr, by ANOVA (A,B, D) or t-test (C).

## Discussion

In this manuscript, we demonstrate for the first time that E2 promotes pro-angiogenic behavior in RVECs from rodents and human patients with RVF. Specifically, in RVECs from rats with RVF as well as from PH patients with RVF, E2 promotes vascular network formation, migration and proliferation and increases abundance of pro-angiogenic mediators. E2 also increases RV vascular density in animal models of PH and RVF. We demonstrate that E2’s effects are ERα- and apelin-dependent and that apelin is downstream of ERα. Selectively activating ERα recapitulates pro-angiogenic effects of E2 and when used in a rescue approach, is sufficient to increase RV vascular density in rats with PH and RVF. Together, these data suggest that activating the E2-ERα-apelin axis may be a novel tool to improve RV capillary density and RV function in PH-induced RVF and other types of RVF.

Elevated RV afterload leads to increased cardiac workload and cardiomyocyte hypertrophy^6,27^. Adequate angiogenesis is critical to maintaining substrate delivery to the hypertrophied myocardium^28,29^. In addition, the endothelium exerts paracrine functions that support the homeostasis and adaptation of the neighboring myocardium^9,27^. While there is an ongoing debate as to whether capillary density is decreased versus maintained in the failing RV, it is an accepted paradigm that RVEC angiogenic function is impaired in the setting of RVF. For example, angiogenic properties of primary cultured RVECs from patients with decompensated RVF in Matrigel tube formation assays are impaired, mediated by decreased miR-126 signaling^7^. Restoring miR-126 function improves RVEC function and RV adaptation^7^. Improving and restoring RVEC angiogenic function therefore is an important therapeutic goal in RVF^4^. While our current as well as several prior reports demonstrate decreased vascular density in RVF^7,30,31^, two other rigorous and well-performed studies questioned whether RV capillary is indeed decreased in RVF^32,33^. Such discrepancies may be due to the study of different animal models and different disease stages as well as differences in analytic methods. However, the proper question may not be whether vascular density is decreased in RVF or not, but rather whether vascular density is *sufficient* for the increase in cardiomyocyte volume and substrate demand of the remodeled heart. For example, RV angiogenesis is an early adaptive response to chronic hypoxia-induced PH^29^. Consequently, an insufficient increase in RV vascular density in the setting of increased workload may lead to uncoupling between demand and supply, thus triggering RV decompensation. It was not the focus of our paper to settle this debate, but rather we sought to determine whether the E2-ERα axis can improve RVEC angiogenic function and increase RV capillary density to augment RV vascular adaptation. In light of prior studies clearly demonstrating impaired angiogenic behavior in RVECs from animals or human subjects with RVF^7,28^, establishing that E2 and ERα restore and augment RVEC angiogenic function therefore is an important and clinically relevant finding.

In systemic vessels, ERα facilitates EC recovery, blocks monocyte adhesion to ECs, and inhibits vasoconstriction^34–40^. Loss-of-function mutations in ERα have been linked to endothelial dysfunction, coronary artery disease, myocardial infarction, and stroke^41–43^. In the LV, ERα, under control of MEF2 and class II HDACs, mediates neo-angiogenesis in the setting of myocardial infarction by upregulating VEGF-A^44^. Pro-angiogenic effects in the setting of myocardial infarction have also been demonstrated for cardiomyocyte ERα via paracrine effects^45^.

ERα effects in the vasculature of the RV-pulmonary artery axis, on the other hand, are poorly defined. Older studies demonstrated stimulatory effects in pulmonary artery ECs (PAECs) on prostacyclin and nitric oxide synthesis^46,47^, and we previously showed that ERα mediates EC-dependent vasorelaxation in isolated pulmonary artery rings^48^. More recently, we demonstrated that ERα is decreased in PAECs from pulmonary artery hypertension (PAH) patients, and that treatment of PAECs with ERα agonist increases expression of the EC-protective mediators BMPR2 and apelin^19^. We also demonstrated a decrease of ERα abundance in RVECs from PAH patients with RVF^17^, suggesting there is a lack of vasculoprotective ERα in this context.

While other groups have demonstrated pro-angiogenic effects of ERβ in the heart^49^, a preponderance of evidence suggest that E2’s pro-angiogenic effects in the heart and blood vessels are primarily mediated via ERα^19,34–40,48^. Our current and prior^17^ data suggest that this paradigm also applies to RVECs.

While ERα has been linked to PH development in prior reports^50,51^, these studies focused on PA smooth muscle cells and hypoxic mice, thus limiting their generalizability. Our previously published data demonstrate that selectively activating ERα improves RV function, attenuates PA remodeling, and improves survival in rats with severe PH, even when used in a rigorous rescue approach^17,19^. Our current data adds new evidence that selectively activating ERα also improves RVEC angiogenic function and RV vascular density. This does not rule out that ERα may have detrimental effects on PA smooth muscle cells in hypoxic mice, but in rat models of severe PH and RVF, as well as in human RVECs, the net effect of ERα seems to be protective.

Sex hormone signaling is complex, and effects often are context-dependent and compartment-specific^18^. Previous approaches aimed at non-selectively enhancing or inhibiting E2 in PH have been conflicting. We posit that such approaches may not be precise enough to therapeutically target this complex and pleiotropic pathway. For example, a phase I trial of aromatase inhibition in PAH^52^, while demonstrating an increase in six-minute walking distance and no change in RV function in the entire population, revealed a decrease in RV function and/or an increase in NT-pro-BNP levels (indicating increased RV stress) in ≥25% of patients^53^. On the other hand, E2 treatment, while improving RV function in experimental PH, may promote PASMC proliferation, and higher E2 levels have been linked to more severe PAH in male and post-menopausal female patients^15,16,51,54^. A recently completed phase II trial of aromatase inhibition in PAH was negative^52^. On the other hand, selectively activating ERα signaling may be a novel and more precise strategy to harness protective estrogenic effects in the RV-PA unit that may maximize benefit while avoiding detriment. This may be a solution to the “estrogen puzzle” in PAH. While the work presented here focuses on protective effects of ERα in ECs and blood vessels in the RV, our prior work identified a cardioprotective ERα-BMPR2-apelin axis in RV cardiomyocytes (RVCMs)^17^, suggesting that ERα’s effects extend beyond angiogenic signaling and may also include pro-contractile effects in RVCMs. Since apelin also exerts anti-inflammatory effects, and given ERα’s inhibitory effects on cells of the immune system^55,56^, it is also conceivable that ERα exerts anti-inflammatory effects in the RV. ERα effects on immune cell function in the RV are currently under investigation in our laboratory.

An ideal RV-targeted therapy in PAH would improve RV function not only by directly by targeting the RV, but also indirectly by reducing RV afterload. Activating ERα signaling could fulfill these criteria, since in addition to targeting RVCMs, ERα agonist treatment also reduces PA remodeling in rats with PH and increases BMPR2 and apelin in PAECs from PAH patients^17,19^, suggesting that activating ERα could be a two-pronged approach that exerts protective effects in both components of the RV-PA axis. Since ERα-selective agonists confer cardiovascular protection without unwanted uterotrophic effects^36^, we suggest that an ERα-targeted strategy is both efficacious and safe and could rapidly be evaluated in clinical trials.

The observation that ERα’s effects are apelin and APLNR-dependent is of particular interest. Apelin is a vasoactive peptide that has recently emerged as an important mediator in PAH development. Via its receptor APLNR, apelin targets ECs, smooth muscle cells and cardiomyocytes^17,57^. Apelin plays a critical role in cardiovascular development as well as LV and lung vascular homeostasis^58,59^. Emerging data suggest a role for apelin in the RV: *Aplnr*^−/-^ mice exhibit dilated and/or deformed RVs, *Apln*^−/-^ mice exhibit exaggerated PH, and circulating apelin levels are decreased in PAH patients^58,60^. In addition, apelin is an inotrope in the RV^61,62^ and is decreased in RVs from rats with SuHx-PH^15^. We previously discovered that E2’s RVCM-protective effects include up-regulation of apelin and that this effect is ERα-dependent^15,17^. We now demonstrate that apelin promotes tube formation in RVECs from SuHx-PH rats and is necessary for E2 and ERα agonist to exert pro-angiogenic effects. Pro-angiogenic effects of apelin in the RV may explain at least in part improved RV function noted in studies of apelin supplementation in PH. In combination with our studies of apelin signaling in RVCMs, our results establish apelin as a critical and indispensable mediator of protective E2-ERα signaling in the RV.

Our study has limitations. First, stereology is considered the gold standard for assessing three dimensional structures. While we employed this technique in proof-of-concept analyses, we did not consistently use stereology due to its limitation of requiring the entire RV to be fixed. This would not allow for isolating RVECs or performing biochemical analyses. Like ours, several important prior studies employed two-dimensional analyses^7,30,31^. In addition, we and others demonstrated that findings of decreased vascular density in vivo are accompanied by decreased tube formation and impaired angiogenic behavior in cultured RVECs^7^, thus supporting the in vivo findings. In addition, we performed stereological quantification in key experimental groups, and the observed results mirrored those of the two-dimensional analysis. We therefore feel confident that two-dimensional analyses, when performed in a random manner and by blinded investigators, is an appropriate technique.

Second, there is an ongoing debate as to whether capillary density indeed is decreased in human RVF^4,7,8,30–33^. It was not the focus of our paper to answer this question, but rather we sought to determine whether the E2-ERα axis is capable of increasing RV capillary density. Availability of human RV tissue is a limitation of the field, and the human tissue employed in one of these studies was not fully characterized and predominantly stemmed from female patients (which are known to have better RV adaptation than their male counterparts). We add to the current literature by demonstrating that E2 can rescue impaired angiogenic function in human RVECs from PH patients. Several studies clearly demonstrated impaired angiogenic behavior in RVECs from animals or human subjects with RVF^8,29^. Impairments in RVEC function and capillarization may also be localized and regional, as recently demonstrated by a study indicating that these processes are localized to fibrotic areas^63^. Establishing that E2 and ERα restore RVEC angiogenic function therefore is an important and clinically relevant finding.

Lastly, while we demonstrate that the E2-ERα axis promotes RVEC angiogenic function in vitro and increases RV capillary density in vivo, it remains unknown whether increasing angiogenesis is necessary for E2 and ERα to improve RV function. Addressing this question is complicated since this would require suppression of angiogenic signaling before or after giving E2 or ERα agonist, which could be associated with confounding effects on other cell types such as cardiomyocytes.

In summary, E2 promotes pro-angiogenic behavior in RVECs from subjects with RVF and increases RV vascular density in vivo in animal models of PAH and RVF. These effects are ERα- and apelin-dependent. Activating the E2-ERα-apelin axis may be a novel tool to improve RV vascular adaptation and RV function in RVF due to PAH as well as other types of PH.

## References

1. Humbert M, Sitbon O, Chaouat A, Bertocchi M, Habib G, Gressin V, Yaici A, Weitzenblum E, Cordier JF, Chabot F, et al. Survival in patients with idiopathic, familial, and anorexigen-associated pulmonary arterial hypertension in the modern management era. Circulation. 2010;122:156–163. doi: 10.1161/CIRCULATIONAHA.109.911818

2. van de Veerdonk MC, Kind T, Marcus JT, Mauritz GJ, Heymans MW, Bogaard HJ, Boonstra A, Marques KM, Westerhof N, Vonk-Noordegraaf A. Progressive right ventricular dysfunction in patients with pulmonary arterial hypertension responding to therapy. Journal of the American College of Cardiology. 2011;58:2511–2519. doi: 10.1016/j.jacc.2011.06.068

3. Vonk Noordegraaf A, Chin KM, Haddad F, Hassoun PM, Hemnes AR, Hopkins SR, Kawut SM, Langleben D, Lumens J, Naeije R. Pathophysiology of the right ventricle and of the pulmonary circulation in pulmonary hypertension: an update. Eur Respir J. 2019;53. doi: 10.1183/13993003.01900-2018

4. Lahm T, Douglas IS, Archer SL, Bogaard HJ, Chesler NC, Haddad F, Hemnes AR, Kawut SM, Kline JA, Kolb TM, et al. Assessment of Right Ventricular Function in the Research Setting: Knowledge Gaps and Pathways Forward. An Official American Thoracic Society Research Statement. Am J Respir Crit Care Med. 2018;198:e15–e43. doi: 10.1164/rccm.201806-1160ST

5. Leopold JA, Kawut SM, Aldred MA, Archer SL, Benza RL, Bristow MR, Brittain EL, Chesler N, DeMan FS, Erzurum SC, et al. Diagnosis and Treatment of Right Heart Failure in Pulmonary Vascular Diseases: A National Heart, Lung, and Blood Institute Workshop. Circ Heart Fail. 2021;14. doi: 10.1161/CIRCHEARTFAILURE.120.007975

6. Vonk Noordegraaf A, Westerhof BE, Westerhof N. The Relationship Between the Right Ventricle and its Load in Pulmonary Hypertension. Journal of the American College of Cardiology. 2017;69:236–243. doi: 10.1016/j.jacc.2016.10.047

7. Potus F, Ruffenach G, Dahou A, Thebault C, Breuils-Bonnet S, Tremblay E, Nadeau V, Paradis R, Graydon C, Wong R, et al. Downregulation of MicroRNA-126 Contributes to the Failing Right Ventricle in Pulmonary Arterial Hypertension. Circulation. 2015;132:932–943. doi: 10.1161/CIRCULATIONAHA.115.016382

8. Frump AL, Bonnet S, de Jesus Perez VA, Lahm T. Emerging role of angiogenesis in adaptive and maladaptive right ventricular remodeling in pulmonary hypertension. Am J Physiol Lung Cell Mol Physiol. 2018;314:L443–L460. doi: 10.1152/ajplung.00374.2017

9. Narmoneva DA, Vukmirovic R, Davis ME, Kamm RD, Lee RT. Endothelial cells promote cardiac myocyte survival and spatial reorganization: implications for cardiac regeneration. Circulation. 2004;110:962–968. doi: 10.1161/01.CIR.0000140667.37070.07

10. Agrawal V, Lahm T, Hansmann G, Hemnes AR. Molecular mechanisms of right ventricular dysfunction in pulmonary arterial hypertension: focus on the coronary vasculature, sex hormones, and glucose/lipid metabolism. Cardiovasc Diagn Ther. 2020;10:1522–1540. doi: 10.21037/cdt-20-404

11. Jacobs W, van de Veerdonk MC, Trip P, de Man F, Heymans MW, Marcus JT, Kawut SM, Bogaard HJ, Boonstra A, Vonk Noordegraaf A. The right ventricle explains sex differences in survival in idiopathic pulmonary arterial hypertension. Chest. 2013. doi: 10.1378/chest.13-1291

12. Ventetuolo CE, Praestgaard A, Palevsky HI, Klinger JR, Halpern SD, Kawut SM. Sex and hemodynamics in pulmonary arterial hypertension. Eur Respir J. 2013. doi: 10.1183/09031936.00027613

13. Ventetuolo CE, Hess E, Austin ED, Baron AE, Klinger JR, Lahm T, Maddox TM, Plomondon ME, Thompson L, Zamanian RT, et al. Sex-based differences in veterans with pulmonary hypertension: Results from the veterans affairs-clinical assessment reporting and tracking database. PLoS One. 2017;12:e0187734. doi: 10.1371/journal.pone.0187734

14. Ventetuolo CE, Ouyang P, Bluemke DA, Tandri H, Barr RG, Bagiella E, Cappola AR, Bristow MR, Johnson C, Kronmal RA, et al. Sex hormones are associated with right ventricular structure and function: The MESA-right ventricle study. Am J Respir Crit Care Med. 2011;183:659–667. doi: 201007-1027OC [pii] 10.1164/rccm.201007-1027OC

15. Frump AL, Goss KN, Vayl A, Albrecht M, Fisher A, Tursunova R, Fierst J, Whitson J, Cucci AR, Brown MB, et al. Estradiol improves right ventricular function in rats with severe angioproliferative pulmonary hypertension: effects of endogenous and exogenous sex hormones. Am J Physiol Lung Cell Mol Physiol. 2015;308:L873–890. doi: 10.1152/ajplung.00006.2015

16. Lahm T, Frump AL, Albrecht ME, Fisher AJ, Cook TG, Jones TJ, Yakubov B, Whitson J, Fuchs RK, Liu A, et al. 17beta-Estradiol mediates superior adaptation of right ventricular function to acute strenuous exercise in female rats with severe pulmonary hypertension. Am J Physiol Lung Cell Mol Physiol. 2016;311:L375–388. doi: 10.1152/ajplung.00132.2016

17. Frump AL, Albrecht M, Yakubov B, Breuils-Bonnet S, Nadeau V, Tremblay E, Potus F, Omura J, Cook T, Fisher A, et al. 17beta-Estradiol and estrogen receptor alpha protect right ventricular function in pulmonary hypertension via BMPR2 and apelin. J Clin Invest. 2021;131. doi: 10.1172/JCI129433

18. Hester J, Ventetuolo C, Lahm T. Sex, Gender, and Sex Hormones in Pulmonary Hypertension and Right Ventricular Failure. Compr Physiol. 2019;10:125–170. doi: 10.1002/cphy.c190011

19. Frump AL, Yakubov B, Walts A, Fisher A, Cook T, Chesler NC, Lahm T. Estrogen Receptor-alpha Exerts Endothelium-Protective Effects and Attenuates Pulmonary Hypertension. Am J Respir Cell Mol Biol. 2023;68:341–344. doi: 10.1165/rcmb.2022-0224LE

20. Rebecca F. Hough CMA, Julie A. Bastarache, Serpil C. Erzurum, Wolfgang M. Kuebler,, Eric P. Schmidt LAS, Steven H. Abman, Diego F. Alvarez, Patrick Belvitch, Jahar Bhattacharya, Konstantin G. Birukov, Stephen Y. Chan, David N. Cornfield, Steven M. Dudek, Joe G. N. Garcia, Elizabeth O. Harrington, Connie C. W. Hsia, Mohammad Naimul Islam, Danny D. Jonigk, Vladimir V. Kalinichenko, Todd M. Kolb, Ji Young Lee, Akiko Mammoto, Dolly Mehta, Sharon Rounds, Jonas C. Schupp, Ciara M. Shaver, Karthik Suresh, Dhananjay T. Tambe, Corey E. Ventetuolo, Mervin C. Yoder, Troy Stevens, and Mahendra Damarla. Studying the Pulmonary Endothelium in Health and Disease An Official American Thoracic Society Workshop Report. American Journal of Respiratory Cell and Molecular Biology. 2024.

21. Harbaum L HJ, Pott J, Ostermann J, Sinning CR, Sau A, Sieliwonczyk E, Ng FS, Rhodes CJ, Tello K, Klose H, Gräf S, Wilkins MR. Sex-Specific Genetic Determinants of Right Ventricular Structure and Function. Am J Respir Crit Care Med. 2024.

22. Provencher S AS, Ramirez FD, Hibbert B, Paulin R, Boucherat O, Lacasse Y, Bonnet S. Standards and Methodological Rigor in Pulmonary Arterial Hypertension Preclinical and Translational Research. Circ Res. 2018.

23. Potus F, Malenfant S, Graydon C, Mainguy V, Tremblay E, Breuils-Bonnet D, Ribeiro F, Porlier A, Maltais F, Bonnet S, Provencher S. Impaired Angiogenesis and Peripheral Muscle Microcirculation Loss Contribute to Exercise Intolerance in Pulmonary Arterial Hypertension. Am J Resp Crit Care Med. 2014;190.

24. Marina Zaitseva DSY, John A Katzenellenbogen, Peter A W Rogers, Caroline E Gargett. Estrogen receptor-alpha agonists promote angiogenesis in human myometrial microvascular endothelial cells. J Soc Gynecol Investig. 2004:529–535. doi: 10.1016/j.jsgi.2004.06.004

25. Emanuela Vitale RR, Marco Lo Iacono, Caterina Cristallini, Claudia Giachino and Raffaella Rastaldo. Apelin-13 Increases Functional Connexin-43 through Autophagy Inhibition via AKT/mTOR Pathway in the Non-Myocytic Cell Population of the Heart. Int J Mol Sci 2022.

26. Frump AL, Albrecht ME, McClintick JN, Lahm T. Estrogen receptor-dependent attenuation of hypoxia-induced changes in the lung genome of pulmonary hypertension rats. Pulm Circ. 2017;7:232–243. doi: 10.1177/2045893217702055

27. Vonk-Noordegraaf A, Haddad F, Chin KM, Forfia PR, Kawut SM, Lumens J, Naeije R, Newman J, Oudiz RJ, Provencher S, et al. Right heart adaptation to pulmonary arterial hypertension: physiology and pathobiology. Journal of the American College of Cardiology. 2013;62:D22–33. doi: 10.1016/j.jacc.2013.10.027

28. Kassa B, Kumar R, Mickael C, Sanders L, Vohwinkel C, Lee MH, Gu S, Poth JM, Stenmark KR, Zhao YY, et al. Endothelial cell PHD2-HIF1alpha-PFKFB3 contributes to right ventricle vascular adaptation in pulmonary hypertension. Am J Physiol Lung Cell Mol Physiol. 2021;321:L675–L685. doi: 10.1152/ajplung.00351.2020

29. Kolb TM, Peabody J, Baddoura P, Fallica J, Mock JR, Singer BD, D’Alessio FR, Damarla M, Damico RL, Hassoun PM. Right Ventricular Angiogenesis is an Early Adaptive Response to Chronic Hypoxia-Induced Pulmonary Hypertension. Microcirculation. 2015;22:724–736. doi: 10.1111/micc.12247

30. Handoko ML, de Man FS, Happe CM, Schalij I, Musters RJ, Westerhof N, Postmus PE, Paulus WJ, van der Laarse WJ, Vonk-Noordegraaf A. Opposite effects of training in rats with stable and progressive pulmonary hypertension. Circulation. 2009;120:42–49. doi: 10.1161/CIRCULATIONAHA.108.829713

31. Bogaard HJ, Natarajan R, Henderson SC, Long CS, Kraskauskas D, Smithson L, Ockaili R, McCord JM, Voelkel NF. Chronic pulmonary artery pressure elevation is insufficient to explain right heart failure. Circulation. 2009;120:1951–1960. doi: 10.1161/CIRCULATIONAHA.109.883843

32. Graham BB, Kumar R, Mickael C, Kassa B, Koyanagi D, Sanders L, Zhang L, Perez M, Hernandez-Saavedra D, Valencia C, et al. Vascular Adaptation of the Right Ventricle in Experimental Pulmonary Hypertension. Am J Respir Cell Mol Biol. 2018;59:479–489. doi: 10.1165/rcmb.2018-0095OC

33. Graham BB, Koyanagi D, Kandasamy B, Tuder RM. Right Ventricle Vasculature in Human Pulmonary Hypertension Assessed by Stereology. Am J Respir Crit Care Med. 2017;196:1075–1077. doi: 10.1164/rccm.201702-0425LE

34. Pare G, Krust A, Karas RH, Dupont S, Aronovitz M, Chambon P, Mendelsohn ME. Estrogen receptor-alpha mediates the protective effects of estrogen against vascular injury. Circulation research. 2002;90:1087–1092.

35. Smirnova NF, Fontaine C, Buscato M, Lupieri A, Vinel A, Valera MC, Guillaume M, Malet N, Foidart JM, Raymond-Letron I, et al. The Activation Function-1 of Estrogen Receptor Alpha Prevents Arterial Neointima Development Through a Direct Effect on Smooth Muscle Cells. Circulation research. 2015;117:770–778. doi: 10.1161/CIRCRESAHA.115.306416

36. Bolego C, Rossoni G, Fadini GP, Vegeto E, Pinna C, Albiero M, Boscaro E, Agostini C, Avogaro A, Gaion RM, et al. Selective estrogen receptor-alpha agonist provides widespread heart and vascular protection with enhanced endothelial progenitor cell mobilization in the absence of uterotrophic action. FASEB J. 2010;24:2262–2272. doi: 10.1096/fj.09-139220

37. Lu Q, Schnitzler GR, Ueda K, Iyer LK, Diomede OI, Andrade T, Karas RH. ER Alpha Rapid Signaling Is Required for Estrogen Induced Proliferation and Migration of Vascular Endothelial Cells. PLoS One. 2016;11:e0152807. doi: 10.1371/journal.pone.0152807

38. Xue B, Pamidimukkala J, Lubahn DB, Hay M. Estrogen receptor-alpha mediates estrogen protection from angiotensin II-induced hypertension in conscious female mice. Am J Physiol Heart Circ Physiol. 2007;292:H1770–1776. doi: 10.1152/ajpheart.01011.2005

39. Billon-Gales A, Fontaine C, Douin-Echinard V, Delpy L, Berges H, Calippe B, Lenfant F, Laurell H, Guery JC, Gourdy P, et al. Endothelial estrogen receptor-alpha plays a crucial role in the atheroprotective action of 17beta-estradiol in low-density lipoprotein receptor-deficient mice. Circulation. 2009;120:2567–2576. doi: 10.1161/CIRCULATIONAHA.109.898445

40. Widder J, Pelzer T, von Poser-Klein C, Hu K, Jazbutyte V, Fritzemeier KH, Hegele-Hartung C, Neyses L, Bauersachs J. Improvement of endothelial dysfunction by selective estrogen receptor-alpha stimulation in ovariectomized SHR. Hypertension. 2003;42:991–996. doi: 10.1161/01.HYP.0000098661.37637.89

41. Schuit SC, Oei HH, Witteman JC, Geurts van Kessel CH, van Meurs JB, Nijhuis RL, van Leeuwen JP, de Jong FH, Zillikens MC, Hofman A, et al. Estrogen receptor alpha gene polymorphisms and risk of myocardial infarction. JAMA. 2004;291:2969–2977. doi: 10.1001/jama.291.24.2969

42. Shearman AM, Cooper JA, Kotwinski PJ, Humphries SE, Mendelsohn ME, Housman DE, Miller GJ. Estrogen receptor alpha gene variation and the risk of stroke. Stroke; a journal of cerebral circulation. 2005;36:2281–2282. doi: 10.1161/01.STR.0000181088.76518.ec

43. Sudhir K, Chou TM, Chatterjee K, Smith EP, Williams TC, Kane JP, Malloy MJ, Korach KS, Rubanyi GM. Premature coronary artery disease associated with a disruptive mutation in the estrogen receptor gene in a man. Circulation. 1997;96:3774–3777.

44. van Rooij E, Fielitz J, Sutherland LB, Thijssen VL, Crijns HJ, Dimaio MJ, Shelton J, De Windt LJ, Hill JA, Olson EN. Myocyte enhancer factor 2 and class II histone deacetylases control a gender-specific pathway of cardioprotection mediated by the estrogen receptor. Circulation research. 2010;106:155–165. doi: 10.1161/CIRCRESAHA.109.207084

45. Mahmoodzadeh S, Leber J, Zhang X, Jaisser F, Messaoudi S, Morano I, Furth PA, Dworatzek E, Regitz-Zagrosek V. Cardiomyocyte-specific Estrogen Receptor Alpha Increases Angiogenesis, Lymphangiogenesis and Reduces Fibrosis in the Female Mouse Heart Post-Myocardial Infarction. J Cell Sci Ther. 2014;5:153. doi: 10.4172/2157-7013.1000153

46. Chen Z, Yuhanna IS, Galcheva-Gargova Z, Karas RH, Mendelsohn ME, Shaul PW. Estrogen receptor alpha mediates the nongenomic activation of endothelial nitric oxide synthase by estrogen. J Clin Invest. 1999;103:401–406.

47. Sherman TS, Chambliss KL, Gibson LL, Pace MC, Mendelsohn ME, Pfister SL, Shaul PW. Estrogen acutely activates prostacyclin synthesis in ovine fetal pulmonary artery endothelium. Am J Respir Cell Mol Biol. 2002;26:610–616.

48. Lahm T, Crisostomo PR, Markel TA, Wang M, Wang Y, Tan J, Meldrum DR. Selective estrogen receptor-{alpha} and estrogen receptor-{beta} agonists rapidly decrease pulmonary artery vasoconstriction by a nitric oxide-dependent mechanism. Am J Physiol Regul Integr Comp Physiol. 2008;295:R1486–1493. doi: 90667.2008 [pii]10.1152/ajpregu.90667.2008

49. Iorga A, Umar S, Ruffenach G, Aryan L, Li J, Sharma S, Motayagheni N, Nadadur RD, Bopassa JC, Eghbali M. Estrogen rescues heart failure through estrogen receptor Beta activation. Biol Sex Differ. 2018;9:48. doi: 10.1186/s13293-018-0206-6

50. Wright AF, Ewart MA, Mair K, Nilsen M, Dempsie Y, Loughlin L, Maclean MR. Oestrogen receptor alpha in pulmonary hypertension. Cardiovasc Res. 2015;106:206–216. doi: 10.1093/cvr/cvv106

51. Mair KM, Wright AF, Duggan N, Rowlands DJ, Hussey MJ, Roberts S, Fullerton J, Nilsen M, Loughlin L, Thomas M, et al. Sex-dependent influence of endogenous estrogen in pulmonary hypertension. Am J Respir Crit Care Med. 2014;190:456–467. doi: 10.1164/rccm.201403-0483OC

52. Kawut SM, Archer-Chicko CL, DiMichele A, Fritz JS, Klinger JR, Ky B, Palevsky HI, Palmisciano AJ, Patel M, Pinder D, et al. Anastrozole in Pulmonary Arterial Hypertension (AIPH): A Randomized, Double-Blind Placebo-Controlled Trial. Am J Respir Crit Care Med. 2016. doi: 10.1164/rccm.201605-1024OC

53. Frump AL LT. Towards Harnessing Sex Steroid Signaling As A Therapeutic Target In Pulmonary Arterial Hypertension (Editorial). Am J Resp Crit Care Med. 2016;in press.

54. Ventetuolo CE, Baird GL, Barr RG, Bluemke DA, Fritz JS, Hill NS, Klinger JR, Lima JA, Ouyang P, Palevsky HI, et al. Higher Estradiol and Lower Dehydroepiandrosterone-Sulfate Levels Are Associated with Pulmonary Arterial Hypertension in Men. Am J Respir Crit Care Med. 2016;193:1168–1175. doi: 10.1164/rccm.201509-1785OC

55. Lippman JMRaME. The role of estrogen receptor signaling in suppressing the immune response to cancer. JCI. 2021.

56. U Islander * MCE, T Chavoshi *, C Jochems *, S Movérare †, S Nilsson ‡, C Ohlsson †, J-Å Gustafsson §, H Carlsten *. Estren-mediated inhibition of T lymphopoiesis is estrogen receptor-independent whereas its suppression of T cell-mediated inflammation is estrogen receptor-dependent. Clin Exp Immunol 2005;139(2):210–215.

57. Dalzell JR, Rocchiccioli JP, Weir RA, Jackson CE, Padmanabhan N, Gardner RS, Petrie MC, McMurray JJ. The Emerging Potential of the Apelin-APJ System in Heart Failure. Journal of cardiac failure. 2015;21:489–498. doi: 10.1016/j.cardfail.2015.03.007

58. Kang Y, Kim J, Anderson JP, Wu J, Gleim SR, Kundu RK, McLean DL, Kim JD, Park H, Jin SW, et al. Apelin-APJ signaling is a critical regulator of endothelial MEF2 activation in cardiovascular development. Circulation research. 2013;113:22–31. doi: 10.1161/CIRCRESAHA.113.301324

59. Kim J, Kang Y, Kojima Y, Lighthouse JK, Hu X, Aldred MA, McLean DL, Park H, Comhair SA, Greif DM, et al. An endothelial apelin-FGF link mediated by miR-424 and miR-503 is disrupted in pulmonary arterial hypertension. Nat Med. 2013;19:74–82. doi: 10.1038/nm.3040

60. Chandra SM, Razavi H, Kim J, Agrawal R, Kundu RK, de Jesus Perez V, Zamanian RT, Quertermous T, Chun HJ. Disruption of the apelin-APJ system worsens hypoxia-induced pulmonary hypertension. Arterioscler Thromb Vasc Biol. 2011;31:814–820. doi: 10.1161/ATVBAHA.110.219980

61. Dai T, Ramirez-Correa G, Gao WD. Apelin increases contractility in failing cardiac muscle. Eur J Pharmacol. 2006;553:222–228. doi: 10.1016/j.ejphar.2006.09.034

62. Brash L, Barnes GD, Brewis MJ, Church AC, Gibbs SJ, Howard L, Jayasekera G, Johnson MK, McGlinchey N, Onorato J, et al. Short-Term Hemodynamic Effects of Apelin in Patients With Pulmonary Arterial Hypertension. JACC Basic Transl Sci. 2018;3:176–186. doi: 10.1016/j.jacbts.2018.01.013

63. Kenzo Ichimura MB, Adam M Andruska, Fan Zhang, Katharina Schimmel, Spencer Bonham, Angela Kabiri, Vitaly O Kheyfets, Shoko Ichimura, Sushma Reddy, Yuqiang Mao, Tianyi Zhang, Gordon X Wang, Everton J Santana, Xuefei Tian, Ilham Essafri, Ryan Vinh, Wen Tian, Mark R Nicolls, Shin Yajima, Yasuhiro Shudo, John W MacArthu, Y Joseph Woo, Ross J Metzger, Edda Spiekerkoetter 3D Imaging Reveals Complex Microvascular Remodeling in the Right Ventricle in Pulmonary Hypertension. Circ Res. 2024. doi: 10.1161/CIRCRESAHA.123.323546

