## Supplementary material for "17β-Estradiol and Estrogen Receptor α Promote Right Ventricle Angiogenesis in Pulmonary Hypertension via Apelin Signaling": Material and methods

### Materials and Methods

All experiments were performed using randomization and blinding at the time of measurement and analysis. We followed published best practices for animal, endothelial cell and sex difference/hormone research[20-22].

*Animal care*

All rodents used in studies were approved by the Indiana University School of Medicine Institutional Animal Care and Use Committee and were adherent with the National Institutes of Health guidelines for care and use of laboratory animals under the animal welfare assurance act. Rats and mice were allowed ad libitum access to food and water and were housed in a facility with a 12-hour light/dark cycle.

*Animal models*

*Sugen/hypoxia-induced pulmonary hypertension (SuHx-PH):* Su5416 (20 mg/kg subcutaneously; dissolved in DMSO; R&D Systems (Tocris; Minneapolis, MN) was administered prior to hypoxia exposure. Animals were housed in a hypobaric hypoxia chamber for 3 weeks followed by a return to room air for 4 weeks as described previously[15,17]. Experimental groups included intact female Sprague-Dawley normoxic controls, intact female SuHx-PH rats, ovariectomized (OVX; described previously[15]) SuHx-PH females, and OVX SuHx-PH females replete with E2 (75 µg/kg/d via subcutaneous pellets; Innovative Research of America; Sarasota, FL)[17]. Females weighed 150-175 g (Charles River; Wilmington, MA) and were 8 weeks old at the time of Su5416 injection. Additional studies were performed in male normoxic Sprague-Dawley rats, SuHx-PH rats, and SuHx-PH rats with E2 (75 µg/kg/day via subcutaneous pellets), ERα-selective agonist 4,4',4''-(4-Propyl-[1H]-pyrazole-1,3,5-triyl)trisphenol (PPT; 850 µg/kg/day via subcutaneous pellets)[17] or ERβ-selective agonist diarylpropionitrile (DPN; 850 µg/kg/day via subcutaneous pellets).[15] Males were 8 weeks old at the time of Su5416 injection and weighed 175-200g. Vehicles for E2, PPT, and DPN were tested in prior studies and found not to have any effects on cardiopulmonary parameters[15].

*Monocrotaline-induced PH (MCT-PH):* Male Sprague-Dawley rats (250-275 g; Charles River) were subcutaneously injected with MCT (60 mg/kg; Sigma Aldrich; St. Louis, MO) dissolved in sterile saline. At 14 days, subgroups of animals received E2 (75 µg/kg/day) or PPT (850 µg/kg/day) for 14 additional days via subcutaneous pellets. Final endpoint was taken at day 28.

Cardiopulmonary hemodynamics, and RV structural, functional and molecular alterations of the animals employed were previously published[15,17].

*Right ventricular endothelial cell (RVEC) isolation and culture*

*Rat RVECs:* RVs were dissected from male normoxic or SuHx-PH Sprague-Dawley rats, minced and digested using Collagenase II (Gibco, ThermoFisher; Waltham, MA) at 37°C for 1-3 hrs as previously described. Cells were selected using Pan-mouse IgG Dynabeads (ThermoFisher) coated with mouse monoclonal anti-rat CD31 antibody (BD Biosciences; BD Biosciences) and then seeded on a gelatin-coated 6-well plate in EGM-2MV (Lonza; Basel, Switzerland) media with charcoal stripped FBS (ThermoFisher) substituting regular FBS and supplemented with Normocin (Invivogen, San Diego, CA). Endothelial lineage was validated based on morphology, endothelial marker expression, matrigel tube formation, and Dio-AC-LDL uptake. Cells were utilized in experiments up to passage 7. To study E2 effects in an E2-naïve background, we focused on cells from male animals.

*Human RVECs:* RVECs from PH patients with RV failure (PH-RVECs) were isolated as previously described[23]. Patient characteristics are provided in **table 1**. Cells were grown in Endothelial Cell Growth Basal Medium-2 supplemented with 0.1% human epidermal growth factor, 0.1% ascorbic acid, 0.1% GA-1000 (gentamicin, amphotericin B), 2% fetal bovine serum, 0.1% VEGF, 0.4% human basic fibroblast growth factor, 0.04% hydrocortisone, 0.1% R3-Insulin-like growth factor -1, and 1% streptomycin-penicillin (all chemicals were purchased from Lonza). Cells were grown at 37°C in a humidified incubator gassed with 5% $CO_2$ and were used for experiments at the third to ninth passage in charcoal stripped media.

*Human cardiac microvascular ECs* (hCMVECs; of male origin) were purchased from Lonza and grown and maintained in EGM-2MV media as described for human RVECs. Cells were utilized in experiments up to passage 9.

*Cell culture and chemical treatment*

Cells were starved overnight in serum and phenol-red free media for at least 12 hours prior to treatment. RVECs were treated with E2 (0.1-100 nM, Sigma Aldrich), the ERα-selective agonists BTPα (100 nM; obtained through academic collaboration with Eli Lilly; Indianapolis, IN)[17] or PPT (10-100 nM; Sigma)[24] or ethanol vehicle control, as well as with apelin receptor antagonist ML221 (100 µM, Tocris)[17] or DMSO vehicle control, or Pyr-Apelin-13 (100 nM, Tocris)[25] or $H_2O$ vehicle control for times indicated.

*siRNA knockdown*

RVECs were transfected at 50% confluency using lipofectamine 2000 or RNAiMax (Thermo Fisher) and Silencer Select siRNA oligos (Thermo Fisher) directed against ERα, ERβ, apelin or scrambled control for 24 hours as directed by the manufacturer. Cells were then treated with E2 at doses and times indicated. Knockdown of target protein was confirmed by Western blot at 48 hours.

*Cell proliferation*

Cell proliferation was evaluated using the CCK8 test (Dojindo, Rockville, MD, USA). $10^4$ cells per well were seeded into 96-well plates overnight and treated with or without E2 (0,1 to 100 nM) for 24 hours in duplicate. Cells were then incubated with 10 µL of CCK8 solution for 3 hours and absorbance was measured at 450 nm using a microplate reader (BioTek, Winooski, VT).

*Tube formation assay*

$5x10^4$ RVECs were plated onto Geltrex LDEV-free phenol red-free reduced growth factor basement membrane matrix (ThermoFisher) or onto Matrigel Growth Factor Reduced Basement Membrane Matrix, Phenol Red-free, LDEV-free (Corning; Corning, NY) in either EBM or EBM-2 basal media for 6 to 12 hours. 15 fields per condition were imaged at 10x magnification using a Nikon Eclipse 80i inverted microscope with camera and NIS-Elements 4.0 software (Nikon Instruments, Melville, NY) and rings were quantified using ImageJ. Experiments were performed in technical triplicate in hCMVECs, PH-RVECs, and in rat RVECs.

*Wound healing assay*

hCMVECs or PH-RVECs (6 x $10^4$) were seeded in 12-well plates. When confluence reached 100%, cells were scratched with a sterile pipette tip to make a gap. Afterward, floating cells and debris were removed by washing with PBS and then treated with or without E2 (0.1-100 nM) for 24 hours in duplicate. Photographic images of the plates were taken under a microscope immediately after healing and every 2 hours for 24 hours. The area of the scratch was measured using ImageJ software.

*Western blotting*

Cells were homogenized and lysed in ice-cold RIPA lysis buffer (Thermo Fisher Scientific) or Cell Lysis Buffer (Cell Signaling) containing proteinase inhibitor cocktail (EMD-Millipore-Sigma Aldrich) and PhosStop inhibitor cocktails (Roche, Pleasanton, CA). After homogenization, lysate was centrifuged, and the supernatant was saved. Protein concentration was measured using BCA Protein Assay (ThermoFisher Scientific). Rabbit anti-VEGF Receptor 2 (Cell Signaling, Danvers, MA), anti-Apelin (LSBio, Lynnwood, WA), anti-Angiopoietin 2 (Invitrogen-ThermoFisher Scientific, Carlsbad, CA) primary antibodies were used at a dilution of 1:1000 and mouse anti-phospho eNOS (Ser 1177) (BD Biosciences), anti-eNOS (BD Biosciences) and anti-beta-actin (Cell Signaling) were used at a dilution of 1:1000 and 1:5000, respectively, all in blocking buffer (Thermo Fisher Scientific). Rabbit-HRP (Azure Biosystems, Dublin, CA, USA) and Mouse-HRP (Azure Biosystems) secondary antibodies were diluted 1:5000 or 1:20000 in blocking buffer. The protein-antibody complexes were detected by an ECL system and quantified by densitometry (ChemiDoc XRS+, Bio-rad, Hercules, CA).

*Lectin staining and capillary density quantification*

Rat RV tissue was formalin-fixed, embedded in paraffin, and cut into 4 μm sections. RV sections were then heated in citrate antigen retrieval buffer (10 mM 126 Sodium Citrate, 0.05% Tween-20, pH 6.0). Sections were co-stained overnight with Isolectin GS-IB4 (1:100, conjugated to Alexa Fluor 488 or 594 as indicated), wheat germ agglutinin (1:2000, conjugated to Alexa Fluor 488 or 594 as indicated) and anti-fade DAPI (Thermo Fisher) mounting media. Capillaries from 15 fields per section from 3-5 rats were quantified. Images were taken using a Nikon Eclipse 80i microscope with camera and NIS-Elements 4.0 software (Nikon Instruments) at 20x magnification.

*RNA-Sequencing*

Bulk RNA-sequencing was performed on RV tissue from female OVX rats with SuHx-induced PH and RV failure in absence and presence of E2 treatment as described previously[26]. The same animals were employed for analysis of RV vascularization. Hemodynamics and RV function in these rats were described previously[15].

*Statistical analyses*

Results are expressed as means±SEM. At least three independent experiments (run in technical duplicates) were performed for all in vitro studies and reported as N. Statistical analyses were performed with GraphPad Prism (La Jolla, CA). Student’s t-test or one-way ANOVA with Tukey’s or Dunnett’s post-hoc correction were used for comparison of experimental groups. Statistically significant difference was accepted at $p<0.05$.
