## Supplementary figures and images for "17β-Estradiol and Estrogen Receptor α Promote Right Ventricle Angiogenesis in Pulmonary Hypertension via Apelin Signaling"

### Suppl figures

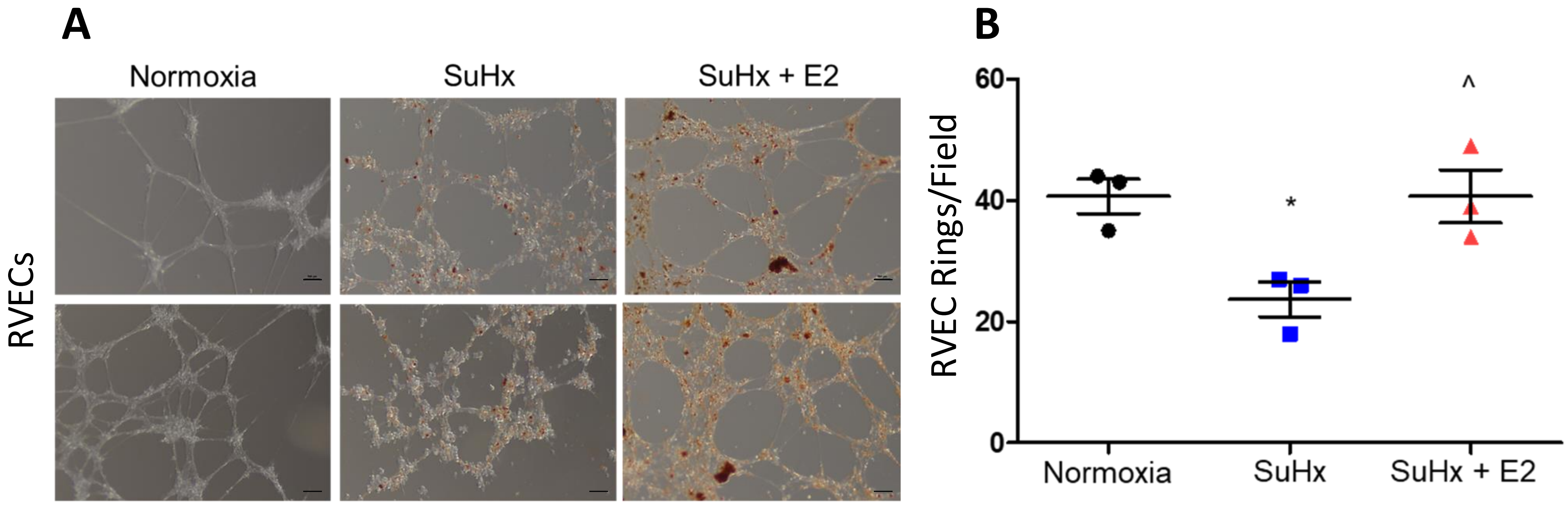

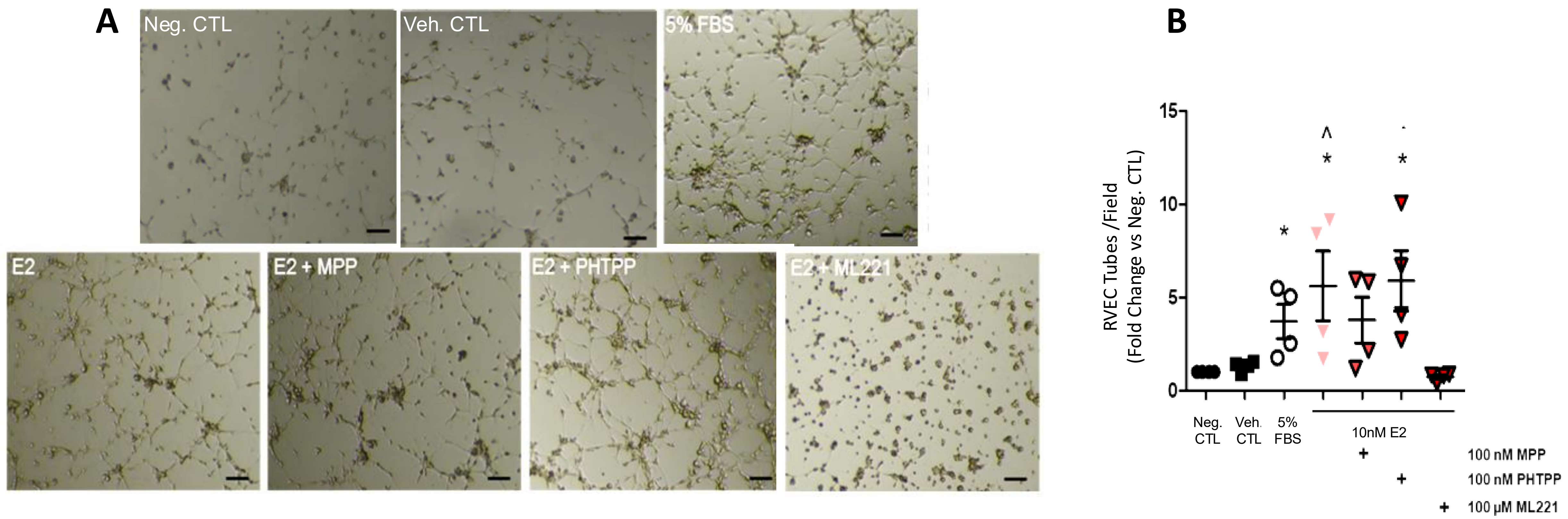

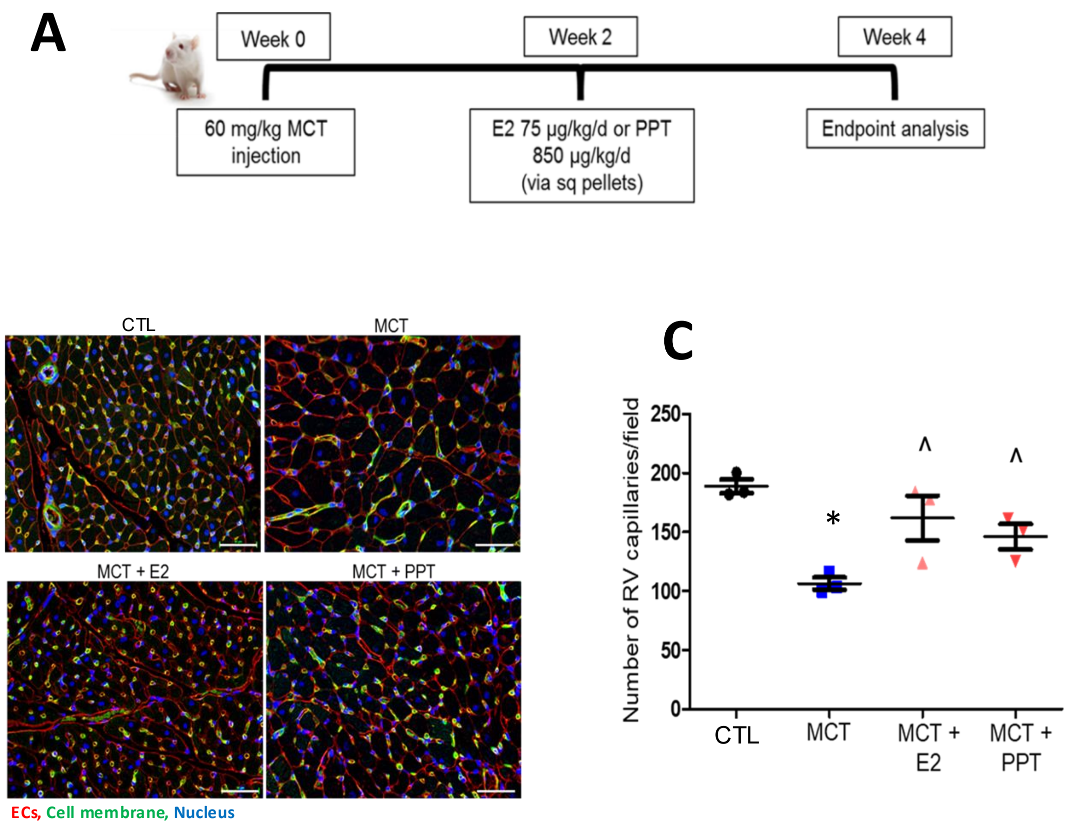
