## Supplementary material for "17β-Estradiol and Estrogen Receptor α Promote Right Ventricle Angiogenesis in Pulmonary Hypertension via Apelin Signaling": Suppl figure legends

**Suppl. Fig. 1: E2 attenuates SuHx-induced impairments in angiogenic function of RV endothelial cells (RVECs). A) and B)** Matrigel tube formation (representative images and quantification) in RVECs from male normoxia control, SuHx-PH or SuHx-PH rats treated with E2 *in vivo*. Images are 4x; scale bar is 100 μm. Brown coloration = CD31-coated dynabeads. Error bars are means±SEM. Each datapoint = one animal. *p<0.05 vs normoxia; ^p<0.05 vs untreated SuHx by ANOVA.

**Suppl. Fig. 2: E2-induced angiogenesis in rat RV endothelial cells (RVECs) is estrogen receptor α (ERα)-dependent. A) and B)** Matrigel tube formation assay using RVECs isolated from control male rats. Images are 4x. MPP = ERα antagonist, PHTPP = ERβ antagonist, and ML221 = apelin receptor antagonist. Concentrations are as indicated in figure; duration of treatment was 24 hours. Error bars are means±SEM. Scale bars=100 μm. *p<0.05 vs neg. CTL; ^p<0.05 vs MPP or ML221 by ANOVA.

**Suppl. Fig. 3: ERα activation with PPT is sufficient to rescue RV vascular rarefaction in male rats with monocrotaline (MCT)-induced PH. A)** Experimental design. **B)** Immunofluorescence overlay of lectin Griffonia simplicifolia (red, endothelial cells (ECs)) wheat germ agglutinin (green, cell membrane) and DAPI (blue, nuclei) in male control, MCT, MCT+E2, & MCT+PPT rats. Images are 20x. Scale bars=50 μm. **C)** Quantification of capillary density as RV capillaries/field. Each datapoint = one rat. Error bars are means±SEM. *p<0.05 vs CTL; ^p<0.05 vs untreated MCT by ANOVA.
